# Chlorine dioxide gas sterilization of cold-chain or temperature-sensitive devices

**DOI:** 10.1101/2025.03.06.641968

**Authors:** Kirk Livingston, Emily Lorcheim, Paul Lorcheim

## Abstract

Cold-chain terminal sterilization is used for products that must be manufactured, stored, and maintained with a carefully controlled temperature range, such as temperature-sensitive drugs and vaccines in prefilled syringes or autoinjectors containing temperature-sensitive drugs. Chlorine dioxide (ClO_2_) is an ideal sterilant for cold-chain terminal sterilization because it forms a true gas at room temperatures (15°C/59°F - 25°C/77°F), with a reliable and consistent process with excellent penetration and does not require elevated temperatures. In the current study, we validated the efficacy of ClO2 on cold stored pre-filled syringes as well as tested ClO_2_-sterilized samples and non-sterilized controls for residual sterilant on the prefilled syringe samples and the ingress of sterilant into the prefilled syringe samples. Residual sterilant remaining on a load after sterilization could cause chemical exposure to patients and negatively affect healthcare worker safety, among other things. Sterilant ingress into a medication could result in chemical contamination, drug degradation, and changes to drug formulation. Few studies have been published on ClO_2_ residue or ingress during terminal sterilization. We demonstrate that chlorine dioxide sterilization leaves residuals and ingress amounts at acceptably low levels. ClO_2_ sterilization is an excellent alternative to ethylene oxide sterilization for cold-chain sterilization because it is sustainable, non-carcinogenic, and does not leave harmful residuals, nor does it ingress into the drug product.

## Introduction

### Sterilization objectives

Effective sterilization renders an object “free from viable microorganisms.” [1-2] Terminal sterilization of medical devices ensures microbial inactivation to a sterility assurance level (SAL) of 10^-6^ for regulatory purposes. [3] Except for terminal sterilization by irradiation, typical sterilization modalities, like ethylene oxide (EtO), require higher temperatures to be effective. High temperatures can compromise temperature-sensitive medical devices, their components, and other medical products that must be stored and transported while maintaining a specific temperature range.

### Cold chain sterilization

“Cold chain” is a set of rules and procedures that help ensure products such as vaccines and medical devices are manufactured, stored, or transported within a controlled temperature range. Cold chain sterilization is a set of parameters requiring that low temperatures be maintained over a controlled range throughout the process.

### Chlorine dioxide sterilization

Chlorine dioxide (ClO_2_) is an ideal sterilization modality for cold chain sterilization because it has high efficacy in penetration and sterilization without elevating temperatures. ClO_2_ sterilization cycles are processed at ambient temperature and typically last 4-6 hours. These cycles do not generate excessive temperatures and keep the product from experiencing damaging temperatures by limiting the product’s time-out-of-refrigeration (TOR). Maintaining temperature parameters is of concern for a significant number of industries. [4] Also, unlike EtO, the devices do not require aeration at elevated temperatures to remove sterilant residues.

In the current study, we tested ClO_2_-sterilized samples and non-sterilized controls for residual sterilant and ingress of sterilant into the prefilled syringe samples. Sterilant ingress into a medication could result in chemical contamination, drug degradation, and changes to drug formulation. Residual sterilant remaining on a load after sterilization could cause chemical exposure to patients and negatively affect healthcare worker safety, among other things. We demonstrate that chlorine dioxide sterilization leaves residuals and ingress amounts at acceptably low levels. We also demonstrate that ClO_2_ is an excellent alternative terminal sterilization choice by virtue of efficacy at ambient temperature parameters and the lack of residue or ingress.

## Materials and methods

### Chlorine dioxide gas

ClO_2_ is a unique single electron-transfer-oxidizing agent known as a stable free radical. ClO_2_ is microbiocidal and is highly soluble in water. Unlike chlorine, its reaction chemistry does not lead to the formation of chlorinated organic products (trihalomethane and chloramine). At use concentrations, ClO_2_ is not flammable, explosive, carcinogenic, or ozone-depleting. After a sterilization cycle, chlorine dioxide gas breaks down into constituent parts, chlorite, chlorate, and chloride, which have relatively low toxicities in humans and are not cytotoxic or mutagenic. [5]

ClO_2_ is a gas above -40°C (-40°F) at sterilization concentrations. Gaseous ClO_2_ is generated with the ClorDiSys CSI CD Cartridge (US EPA label # 80802-1) and is registered with the US Environmental Protection Association as a sterilant. [6] The US-EPA label specifies that chlorine dioxide can be used for sterilization applications. Prefilled syringes and medical devices are typically sterilized in an ambient temperature vacuum sterilizer.

## Materials

- 1800 prefilled syringes (PFS) (Type 1 HPLC Grade distilled water) prepared under controlled conditions to be used as a maximum load
- 9 internal process challenge devices (iPCDs) with biological indicators (*Geobacillus stearothermophilus*) placed inside (**Error! Reference source not found.**). iPCDs by ClorDiSys (New Jersey, USA).
- 9 external process challenge devices (ePCDs) with biological indicators (*Geobacillus stearothermophilus*) placed inside. The housing was produced by ClorDiSys.
- 11 humidity and temperature monitors (TempTale/Sensitech—Massachusetts, USA)
- Sterilization chambers (Steridox-100 vacuum pressure chambers, produced by ClorDiSys)

### Cold-chain parameters for PFS

Restrictions on the handling PFS included maintaining the product between 2°C and 8°C (35.6°F–46.6°F). These restrictions are typical of the handling requirements of cold-chain products.

### Sterilization process

The Steridox-100™ ClO_2_ sterilizer was used to sterilize the PFS. The sterilization process typically consists of a series of steps using controlled time, temperature, and pressure conditions.

### ClO_2_ validation process

Validation is a standard part of every sterilization process and demonstrates its effectiveness and reliability. Validation of a sterilization load (for example, a medical device or combination product) using chlorine dioxide gas sterilization falls under the guidance of ISO 14937:2009. [7] Validation of a sterilization process can be accomplished in several ways. Every validation method must demonstrate that the natural bioburden on a load is killed to a SAL of 10^-6^ during the process. The validation method used in this experiment was the overkill method.

### The overkill method for sterilization validation

The overkill method is a conservative approach to terminal sterilization that goes beyond the minimum requirements needed to kill the target microorganisms, resulting in a high level of sterility assurance. The method uses a standardized, highly resistant microbial spore, typically *Geobacillus stearothermophlfus or Bacillus atrophaeus*, as a biological indicator (BI). Both are more challenging to kill than the natural bioburden found on the load to be sterilized. [8] In this project, *Geobacillus stearothermophlfus* was utilized. After exposure to sterilization, BIs are incubated in a growth-promoting medium under specific conditions to check for the growth of viable spores. No growth (a negative result) indicates the sterilization cycle was effective. The overkill method uses changes in process variables and varying load quantities to establish a sterilization window of minimum and maximum levels of sterilization.

### Biological Indicators demonstrate successful kill

Choosing a sublethal dose of sterilant is often the first step when determining the hierarchy of exposure levels. Exposure levels range from sublethal to lethal to overkill. The sublethal cycle, purposefully set below the expected lethal limit, used suboptimal sterilization parameters to kill some but not all the spores on the BIs, in particular, the ePCD. This nonlethal level becomes the known lower limit for subsequent sterilization cycles. The purpose of the sublethal cycle is to confirm that the ePCD is the “worst case” when compared to the iPCD and the natural bioburden on the load. All subsequent sterilization cycles will incorporate process variables above the lower limit.

The validation process is accomplished by placing BIs within the load in the sterilization chamber (iPCDs) (Figure 3) and alongside the load (ePCDs) (Figure 2), including difficult-to-reach locations (Figure 3). ePCDs are placed externally to the load and must be more challenging to kill than the natural bioburden and the iPCD within the load. The validation process continues by performing sterilization cycles at sublethal levels and then at lethal levels. After each cycle, the BIs are collected and processed to ensure spores are killed to the required SAL of 10^-6^. Ongoing sterilization cycles are then set at the lethal limit, with a half-cycle being 50% of the dosage of chlorine dioxide as measured in ppm/hours.

**Figure 1.**
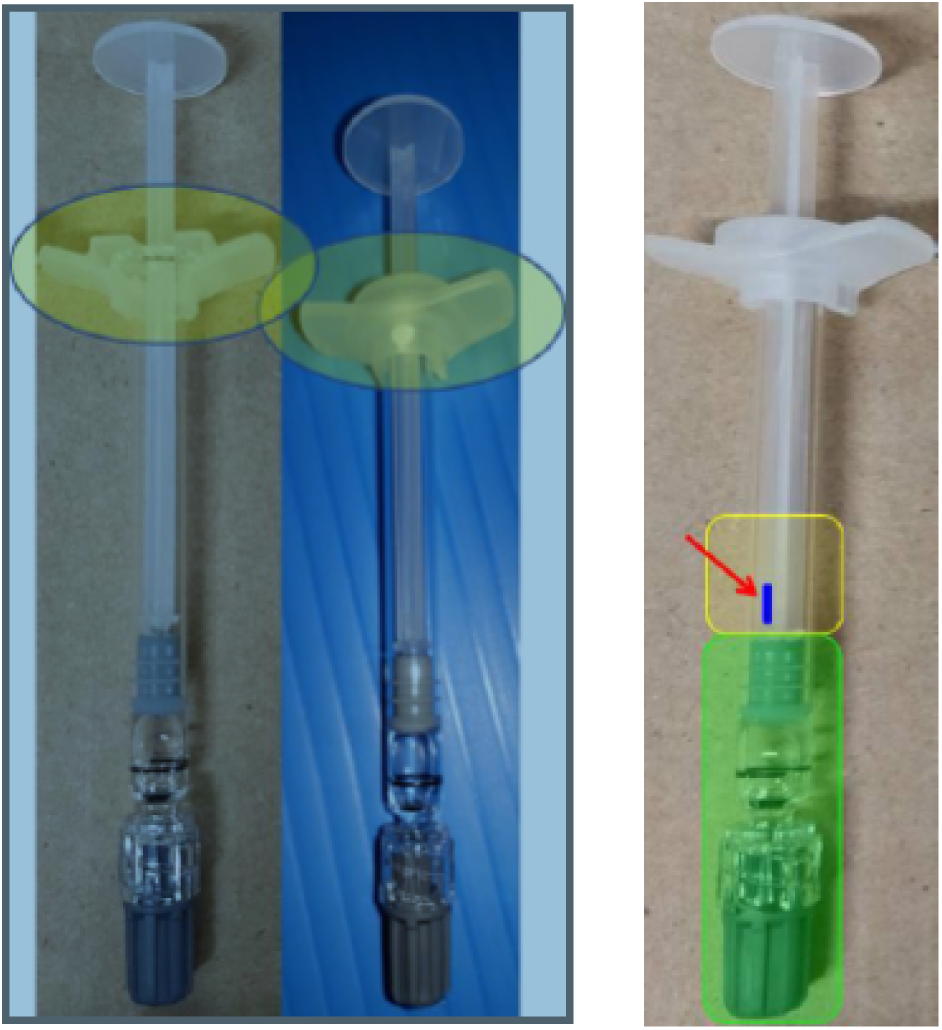
iPCD: Biological indicator Placed Within the Load (a prefilled syringe).

**Figure 2.**
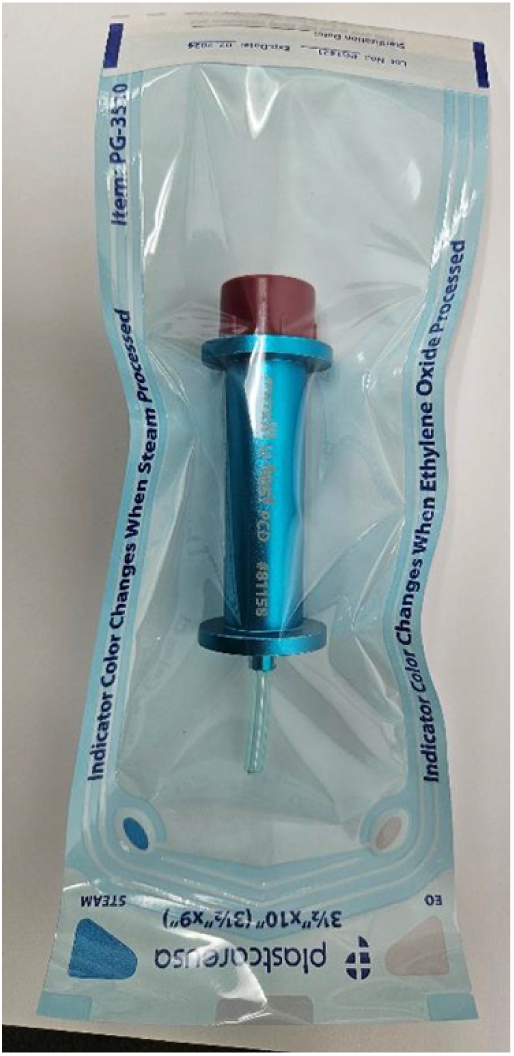
ePCD Within Sterilization Pouch, Which is Placed Alongside the Load in the Sterilization Chamber.

**Figure 3.**
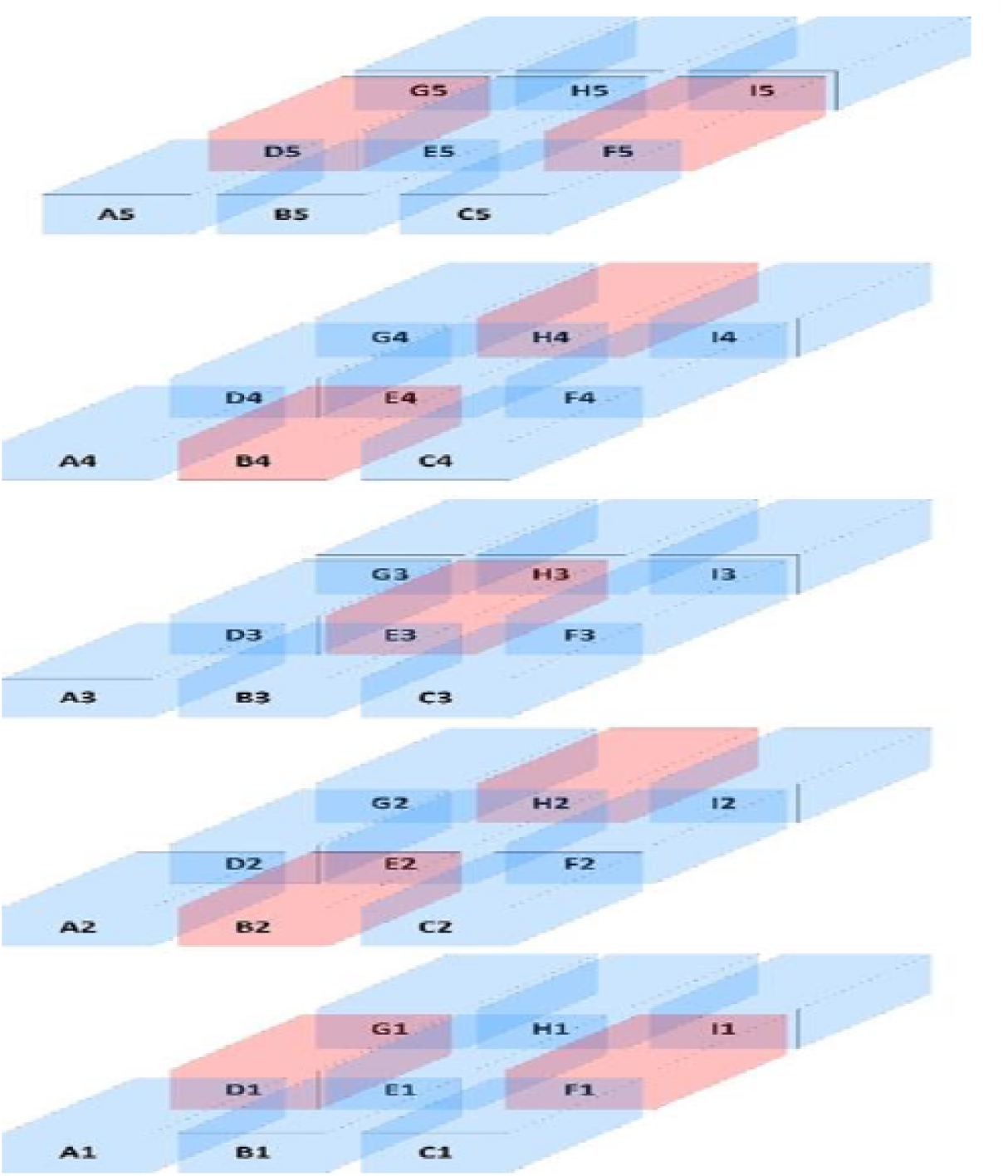
Product Layout in the Sterilization Chamber. Red Designates Locations of iPCDs, ePCDs, and Dataloggers.

### Our validation process included 19 cycles

Our validation process used nineteen sterilization cycles to test a variety of load sizes and cycle parameters. Cycles included:

- One sublethal/fractional exposure cycle to establish the lethality hierarchy between the load’s natural bioburden and the iPCDs and ePCDs.
- Nine half-cycles at a variety of load sizes. The half-cycles used the cycle parameters needed to achieve lethality on the natural bioburden and a full kill on the iPCDs and ePCDs.
- Three full and half-cycles at minimal load
- One full and half-cycle at intermediate load

All loads were processed per protocol, except for the first load which was a sublethal/fractional load. Each load was processed using both a half and a full sterilization cycle. Each half cycle exposed the product to approximately 50% of the dosage of chlorine dioxide as measured in ppm/hour. In addition to the half and full sterilization cycles, the loads consisted of maximum, minimum, and an intermediate quantity of PFS.

The validated chlorine dioxide cycle was performed at ambient temperature, maintaining the cold-chain parameter by avoiding temperatures above 21°C. The average cycle lasted three hours and forty-five minutes, including preconditioning and aeration.

### Residual and ingress testing

Residual tests are designed to demonstrate the absence or quantifiable presence of chlorite, chloride, and chlorate on the outside of the PFS after ClO_2_ sterilization. Ingress tests are designed to demonstrate the absence or quantifiable presence of chlorite, chloride, and chlorate via gas permeation or leakage within the cap, syringe barrel, plunger stopper, or liquid during ClO_2_ sterilization.

After a cycle, the gas rapidly degrades to its constituent parts. Therefore, no chlorine dioxide was expected to remain as a residual (whether as a liquid or a solid) on the outside of the PFS or the inside (cap, syringe barrel, plunger stopper, or liquid) during ingress testing. Instead, the byproducts of chlorite, chloride, and chlorate were tested for residual ingress on the PFS.

Baron Analytical Laboratories (Connecticut, USA) performed the residual and ingress tests per its validated ion chromatography process, which can detect chloride, chlorite, and chlorate levels to parts per million. Ion chromatography is an established and well-used testing method that separates ionized molecules based on differences in charge properties. [9-10]

### Evaluation criteria for chlorite and chlorate levels

Currently, there is no harmonized standard or common specification for ClO_2_ residual levels on medical devices like PFS. However, toxicity guidelines are in place for the amount of chlorate and chlorite in drinking water (Table 1). According to the EPA, chlorite’s maximum contaminant level goal is 0.8 mg/L, and the maximum containment level for chlorite is 1.0 mg/L. [11] Although chlorate is not included in the EPA guidance, the World Health Organization has published a provisional guideline of 0.7 mg/L for chlorate amounts in drinking water. [12]

**Table 1.** EPA Acceptable Levels of Chlorite and Chlorate in Drinking Water.

| Chemical Tested | Maximum Contaminant Level<br>Goal (mg/L) | Maximum Contaminant Level<br>(mg/L) |
| --- | --- | --- |
| Chlorite | 0.8 | 1.0 |
| Chlorate | 0.7 | N/A |

Drinking water guidelines serve as conservative parameters for measuring the toxicity of chlorite and chlorate on devices since many devices will not be ingested.

Although chloride results are included in the testing, the chloride levels are not included in the toxicity analysis. Chloride ions are abundant elements responsible for ionic homeostasis, osmotic pressure, and acid-based balance in the human body. [13] According to the World Health Organization, chloride concentrations in water exceeding 250 mg/L may affect the taste of the water. [14] Chloride is not considered toxic unless levels in the body are unbalanced. [15]

### Testing methods

#### Preparation summary

For ingress testing, one hundred forty (140) syringe components were assembled and filled with type 1 high-performance liquid chromatography (HPLC) grade distilled water to the fill line of l70 μL. Seventy (70) filled syringes were used as blanks or controls, and the remaining 70 were exposed to chlorine dioxide sterilization.

#### Sample preparation

Samples included 200 syringes: 140 syringes filled with 170 µL of type 1 HPLC grade distilled water and 60 unfilled syringes. Seventy filled syringes were exposed to ClO_2_ sterilization, and 70 were used as controls. Thirty unfilled syringes were exposed to ClO_2_ sterilization, and 30 were used as controls.

Filled samples were split into three groups of 23 syringes (with one group having 24). Syringe contents of each group were composited into 4 mL glass vials, which scientists then ran in duplicate for the three separate analytes (chlorite, chlorate, chloride). Unfilled samples were split into three groups of 10 syringes. Each group was composited in 250 mL of deionized water for two hours at room temperature. Solutions from each composite were run in duplicate for the three separate analytes.

#### Testing for sterilant ingress

Each set of seventy syringes was then transferred to standard lab beakers for ion chromatography. Each set was averaged to get the final amount of chlorite, chlorate, and chloride in each sample.

#### Testing for residuals

For residual testing, sixty (60) syringe components were assembled. Thirty (30) assembled and unfilled syringes were used as blanks or controls, and the remaining 30 were exposed to ClO_2_ sterilization.

To remove the residuals from the PFS, we submerged the PFS in a known amount of solution per ISO 11993-7. [16] A verification test at Baron Labs demonstrated that almost all of the residue was dissolved by the solution prior to the ion chromatography testing concentrations using EPA Method 300.0, Determination of Inorganic Anions by Ion Chromatography. [17]

## Results

### Residual testing results

Residual testing results showed that although the amount of chlorite was slightly above the contaminant level goal of 0.8 mg/L, the level was well below the maximum contaminant level of 1.0 mg/L. Table 2 lists a summary of results from residual testing, which demonstrates that residuals were within acceptable limits.

**Table 2.** Residual Testing Summary Table.

|  | Sample Prefill Syringes (PFS) |  |
| --- | --- | --- |
|  | PFS control (unsterilized) | PFS--Sterilized |
| **Average Chlorite** | 0.74 mg/L | 0.82 mg/L |
| **Maximum Contaminant Level Goal** | <0.8 mg/L |  |
| **Maximum Contaminant Acceptable Level** | 1.0 mg/L |  |
| **Result (Pass/Fail)** |  | Pass |
| **Average Chloride** | 1.36 mg/L | 1.13 mg/L |
| **Acceptable Limit** | N/A |  |
| **Result (Pass/Fail)** |  | Pass |

PFS: prefilled syringe; avg: average.

### Ingress testing results

Ingress testing demonstrated that chlorite, chlorate, and chloride levels were below both the maximum contaminant level goal and the maximum contaminant level (Table 3).

**Table 3.**
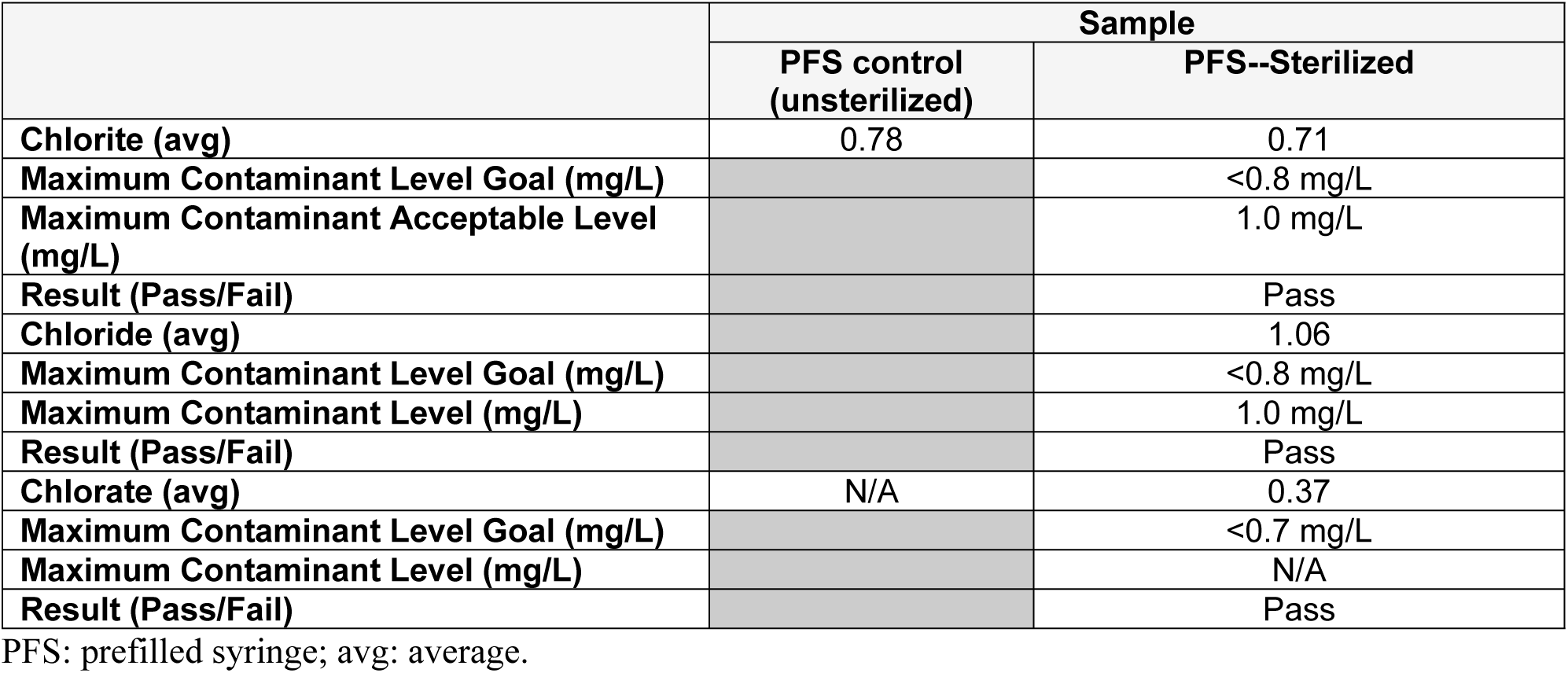
Ingress Testing Summary Table.

## Discussion

Residue testing demonstrated that the presence of chlorite, chloride, and chlorate were within acceptable limits. Ingress testing demonstrated that any chlorite, chloride, and chlorate that had permeated or leaked with the cap, syringe barrel, plunger stopper, or liquid during sterilization were within acceptable limits.

Medical device manufacturers seek alternatives to EtO terminal sterilization, given the global concerns about its sustainability. [18] Chlorine dioxide presents as a viable alternative to EtO sterilization, especially in cold-chain sterilization.

In this experiment, water served as a surrogate for the protein. The validated chlorine dioxide cycle was performed at ambient temperature, allowing the product to avoid experiencing temperatures over 21°C. The average cycle lasted three hours and forty-five minutes, including preconditioning and aeration. In comparison, the ClO_2_ sterilization process kept temperatures within the acceptable range for cold-chain sterilization so the drug would not degrade. Table 4 compares the EtO sterilization with the ClO_2_ sterilization process.

**Table 4.** Comparison: EtO vs. ClO_2_ Sterilization Process.

| Step | Temperature Range | Time Range |
| --- | --- | --- |
| **Ethylene Oxide Full Cycle** |  |  |
| Preconditioning | 38-49°C | 18-24 hours |
| Conditioning | 31-42°C | 59-79 minutes |
| Gas Exposure | 31-42°C | 2.75-3 hours |
| Heated Aeration | 38-55°C | 12-18 hours |
| **Chlorine Dioxide Full Cycle** |  |  |
| Preconditioning | 21°C | 12 minutes |
| Conditioning | 20°C | 60 minutes |
| Gas Exposure | 21°C | 2 hours |
| Aeration | 20°C | 30 minutes |

### Advantages of chlorine dioxide sterilization

ClO_2_ is an effective choice as a terminal sterilant for cold chain sterilization needs. In addition to avoiding temperature spikes and minimizing TOR, ClO_2_ sterilization minimizes cycle length, suggesting that supply chain timing can also be shortened. ClO_2_ also provides the capacity for in-house sterilization by the manufacturer. The ClO_2_ sterilization process is also much simpler than other alternative sterilization modalities because of the ability to sterilize in unit cartons, with instructions for use (IFUs), and other packaging. This streamlines the process and avoids additional logistical complexities. ClO_2_also eases environmental and health concerns compared to EtO and other sterilization methodologies.

### Limitations

This experiment describes ingress and residual testing on syringes prefilled with type 1 HPLC grade distilled water, which may not be generalizable to all medications. Further testing is needed to determine if ClO_2_ sterilization affects different types of medications. Future experiments could use the same method to test ingress and residuals on other types of medical devices, such as inhalers or devices to treat corneal abrasions.

## Conclusion

The sterilization of the PFS using ClO_2_ effectively maintained the required temperatures and kept residual and ingress chlorite, chloride, and chlorate within acceptable levels.

###

## Appendix—detailed testing results

### Residual testing detailed results

Table 5 provides detailed information for residual testing.

**Table 5.** Residual Testing Detail.

| Sample ID | Testing Date | Retention Time (Min) | Amount (mg/L) | Peak Area (UNITS) | Peak Height (UNITS) | Ratio (Area to Height) |
| --- | --- | --- | --- | --- | --- | --- |
| 176458-2-1—Control/Blanks (Not exposed to chlorine dioxide gas for sterilization) |  |  |  |  |  |  |
| Chlorite | 5/26/22 | 3.83 | 0.76 | 63629 | 2622 |  |
| Chloride | 5/26/22 | 4.93 | 1.56 | 505280 | 21106 |  |
| 176458-2-2—Control/Blanks (Not exposed to chlorine dioxide gas for sterilization) |  |  |  |  |  |  |
| Chlorite | 5/26/22 | 3.82 | 0.70 | 53162 | 2265 |  |
| Chloride | 5/26/22 | 4.97 | 1.17 | 13324 | 510 |  |
| 176458-2-3—Control/Blanks (Not exposed to chlorine dioxide gas for sterilization) |  |  |  |  |  |  |
| Chlorite | 5/26/22 | 3.83 | 0.76 | 63069 | 2641 |  |
| Chloride | 5/26/22 | 4.95 | 1.35 | 432708 | 17057 |  |
| 176458-4-1—Exposed to chlorine dioxide gas for sterilization |  |  |  |  |  |  |
| Chlorite | 5/26/22 | 3.85 | 0.84 | 75791 | 3151 |  |
| Chloride | 5/26/22 | 4.92 | 1.09 | 341800 | 12649 |  |
| 176458-4-2—Exposed to chlorine dioxide gas for sterilization |  |  |  |  |  |  |
| Chlorite | 5/26/22 | 3.83 | 0.84 | 75342 | 3268 |  |

| Sample ID | Testing<br>Date | Retention<br>Time<br>(Min) | Amount<br>(mg/L) | Peak<br>Area<br>(UNITS) | Peak<br>Height<br>(UNITS) | Ratio<br>(Area to<br>Height) |
| --- | --- | --- | --- | --- | --- | --- |
| Chloride | 5/26/22 | 4.92 | 1.46 | 469400 | 17848 |  |
| 176458-4-3—Exposed to chlorine dioxide gas for sterilization |  |  |  |  |  |  |
| Chlorite | 5/26/22 | 3.82 | 0.79 | 67934 | 3216 |  |
| Chloride | 5/26/22 | 4.87 | 0.83 | 249039 | 10782 |  |

### Ingress testing detailed results

Table 6 provides detailed information for ingress testing.

**Table 6.**
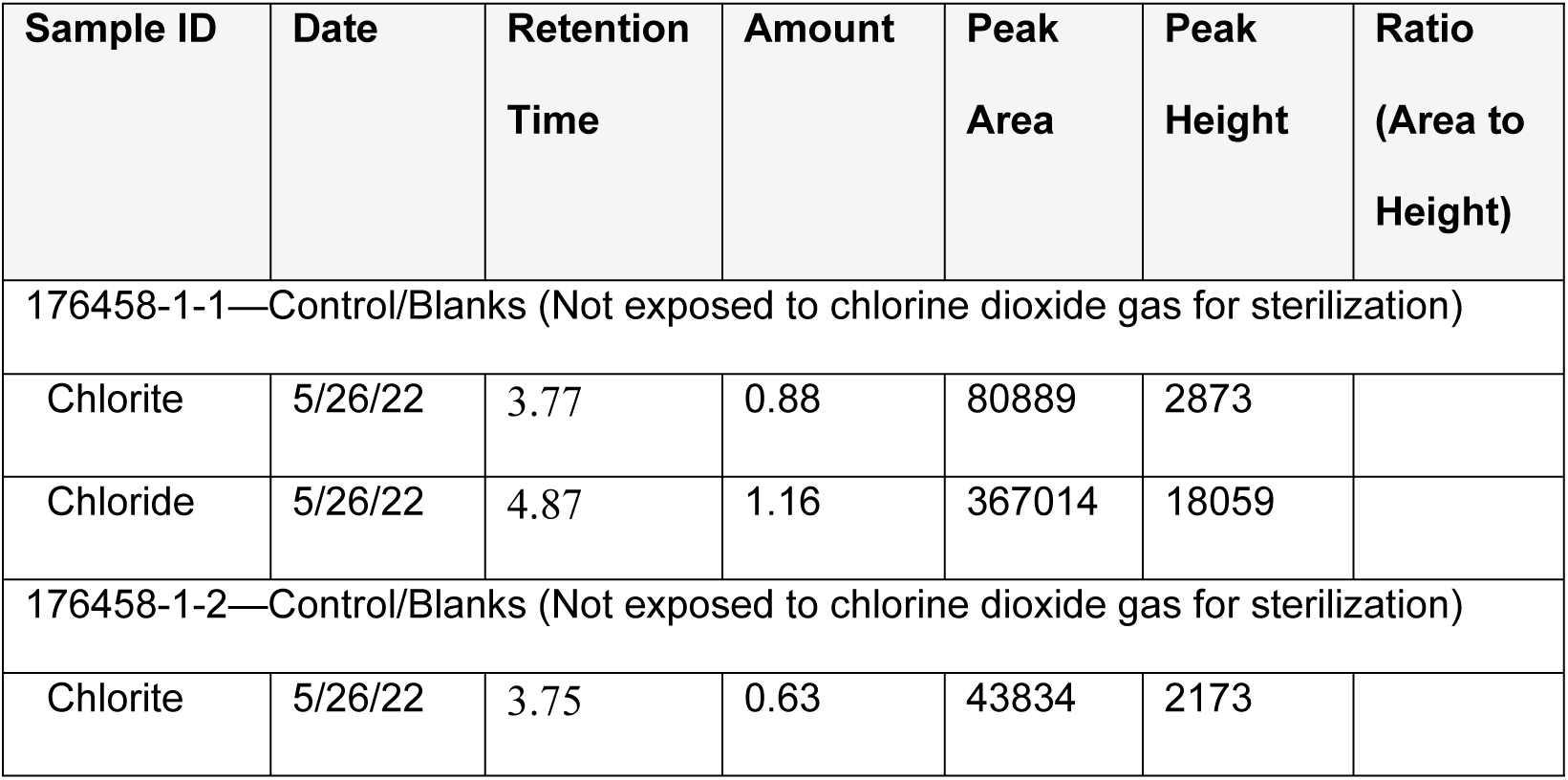
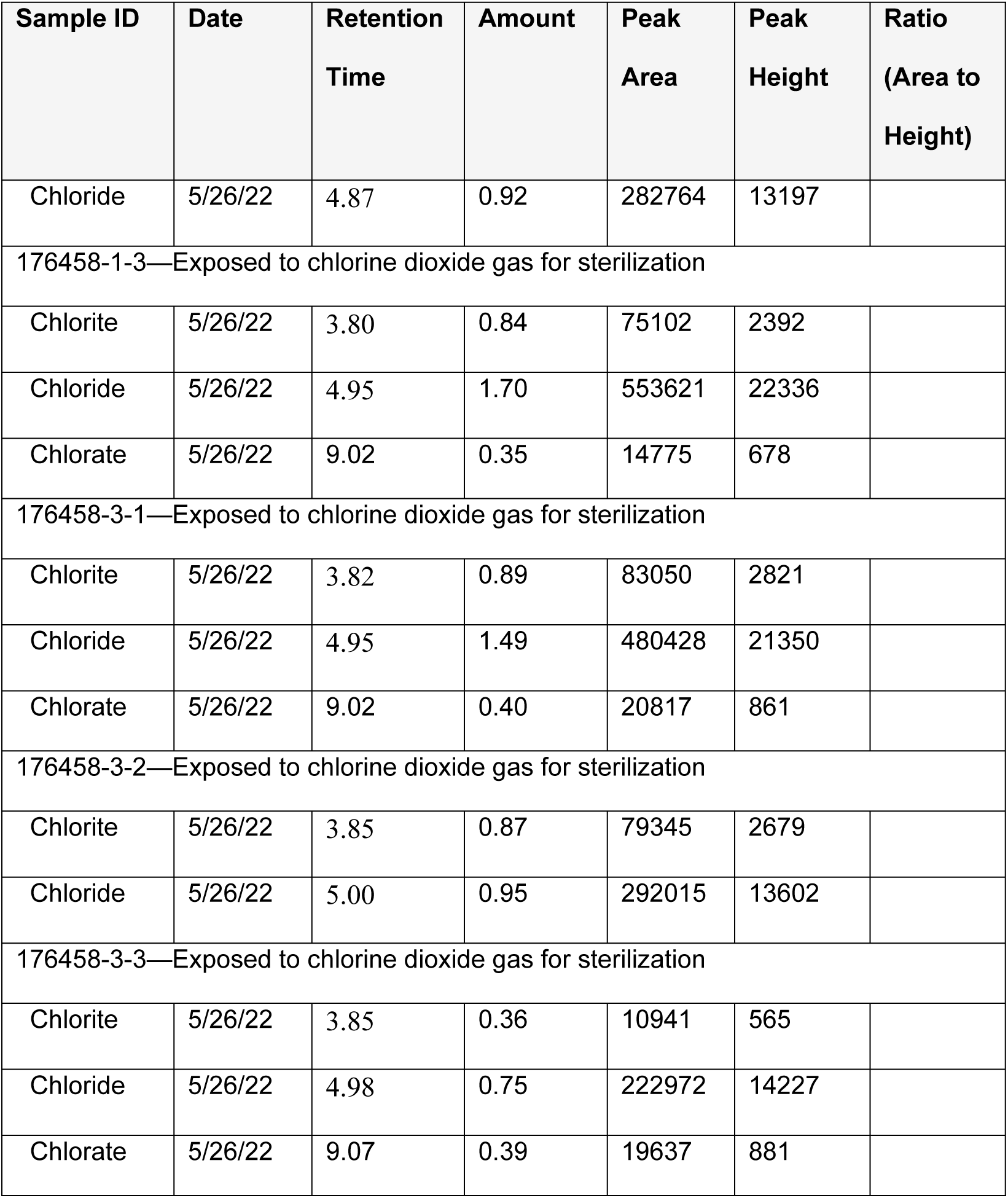
Ingress Testing Detail.

| Sample ID | Date | Retention<br>Time | Amount | Peak<br>Area | Peak<br>Height | Ratio<br>(Area to<br>Height) |
| --- | --- | --- | --- | --- | --- | --- |
| 176458-1-1—Control/Blanks (Not exposed to chlorine dioxide gas for sterilization) |  |  |  |  |  |  |
| Chlorite | 5/26/22 | 3.77 | 0.88 | 80889 | 2873 |  |
| Chloride | 5/26/22 | 4.87 | 1.16 | 367014 | 18059 |  |
| 176458-1-2—Control/Blanks (Not exposed to chlorine dioxide gas for sterilization) |  |  |  |  |  |  |
| Chlorite | 5/26/22 | 3.75 | 0.63 | 43834 | 2173 |  |
| Chloride | 5/26/22 | 4.87 | 0.92 | 282764 | 13197 |  |
| 176458-1-3—Exposed to chlorine dioxide gas for sterilization |  |  |  |  |  |  |
| Chlorite | 5/26/22 | 3.80 | 0.84 | 75102 | 2392 |  |
| Chloride | 5/26/22 | 4.95 | 1.70 | 553621 | 22336 |  |
| Chlorate | 5/26/22 | 9.02 | 0.35 | 14775 | 678 |  |
| 176458-3-1—Exposed to chlorine dioxide gas for sterilization |  |  |  |  |  |  |
| Chlorite | 5/26/22 | 3.82 | 0.89 | 83050 | 2821 |  |
| Chloride | 5/26/22 | 4.95 | 1.49 | 480428 | 21350 |  |
| Chlorate | 5/26/22 | 9.02 | 0.40 | 20817 | 861 |  |
| 176458-3-2—Exposed to chlorine dioxide gas for sterilization |  |  |  |  |  |  |
| Chlorite | 5/26/22 | 3.85 | 0.87 | 79345 | 2679 |  |
| Chloride | 5/26/22 | 5.00 | 0.95 | 292015 | 13602 |  |
| 176458-3-3—Exposed to chlorine dioxide gas for sterilization |  |  |  |  |  |  |
| Chlorite | 5/26/22 | 3.85 | 0.36 | 10941 | 565 |  |
| Chloride | 5/26/22 | 4.98 | 0.75 | 222972 | 14227 |  |
| Chlorate | 5/26/22 | 9.07 | 0.39 | 19637 | 881 |  |

